# NEK7 activates the NLRP1 Inflammasome

**DOI:** 10.1101/2022.12.21.521400

**Authors:** Inés Muela-Zarzuela, Andrea Gallardo-Orihuela, Almudena Pino-Ángeles, Juan Miguel Suarez-Rivero, Daniel Boy-Ruiz, Marta de Gregorio-Procopio, Javier Oroz, Gabriel Mbalaviele, Mario D. Cordero

## Abstract

Inflammasomes including those assembled by NLRP1 and NLRP3 regulate the innate immune system by inducing interleukin (IL)-1β and IL-18 maturation. Inflammasomes are functionally regulated by post-translational modifications such as phosphorylation. The current paradigm posits that NEK7 is the essential and seletive activator of NLRP3; whether this kinase interacts with NLRP3 structurally-related member, NLRP1, has never been explored. Here, we find that NEK7 binds to NLRP1 and promotes its activation independently of NLRP3. IL-1β maturation induced by NLRP1 or NLRP3 inflammasome activators, but not those of the NLRC4 or AIM2 inflammasome is impared in Nek7 deficient cells. This discovery expands the spectrum of NEK7 actions in the regulation of inflammasome pathways.

## Introduction

NLRP (NLR, Pyrin domain containing domain) family members share a conserved tertiary structure comprising an N-terminal pyrin domain (PYD), a central nucleotide-binding and oligomerization domain (NATCH) and a variable number of C-terminal leucine-rich repeats (LRR) (Wang et al., 2021). NLRP1 LRR has an extra function to find (FIIND) domain with autolytic activity followed by a CARD domain (Hollingsworth et al., 2021).

NEK7 is a small protein that binds to the LRR and NATCH domains of NLRP3 and promotes its canonical activity (Sharif et al., 2019). LRR is required for the inactive NLRP3 cage-like structure (Wang et al., 2021), but its function in other NLR proteins remains unclear and it is not present in other receptors such as NLRP10 (Damm et al., 2013). NLRP1 and NLRP3 inflammasomes share a set of common activators. For instance, both inflammasomes are activated in response to several pathogen-associated molecular patterns (PAMPs) and danger-associated molecular patterns (DAMPs) and are concomittantly involved in a wide range of human diseases (de Zoete et al., 2014, Tupik et al., 2020).

The recently reported Cryo-EM structure of NEK7 bound to the inactive and active forms of NLRP3 (Sharif et al., 2019, Xiao et al., 2022) open up the possibility of exploring the interaction of this kinase with other NLRs structurally similar to NLRP3. To this end, we performed a structural modeling of NLRP1-NEK7 complex. Specifically, we examined and compared pairwise interactions involved in the binding of NEK7 with NLRP1 or NLRP3 and tested their dynamics and stability by means of all-atom Molecular Dynamics (MD) simulations alongside with biochemical studies. The present study identified NEK7 as a critical protein that activates the NLRP1 inflammasome.

## Results and Discussion

Inflammasomes share a similar spatial configuration of their functional domains based on current available structures (Wang et al., 2021). As a result, it is reasonable to predict that key inflammasome processes such as priming, activation, or oligomerization might follow these structural features. Therefore, we analyzed specific pairwise interactions between NEK7 and the NLRP1 or NLRP3, and their evolution through MD trajectories as observed in the cryo-EM structure of NLPR3-NEK7 (Sharif et al., 2019). The residues in the LRR domain and the Helical Domain 2 (HD2) of NLRP3 that interact with NEK7 were also conserved in NLRP1 domains. Among the ones described in the cryo-EM structure, R131 in NEK7 interacted with D747 in most of the replicas (Fig. 1A), whereas other residues with drastic effects on the complex, such as Q129 and R136, seldom engaged into long-lasting interactions with NLRP3. Adjacent residues, including R121 or K128, that were not observed to interact in the cryo-EM structure, formed hydrogen bonds with residues D807 and E864 in the LRR domain of NLRP3 in our trajectories. With regard to NLRP1, R121, Q129 and R131 interacted with multiple charged (E717, E740, E812), polar (S761, Q900, Q931) and even aromatic residues (F874) in the LRR domain. This extensive network also included positively charged residues in NEK7 (K124, K128, K140) interacting with multiple different residues in NLRP1. Other interface areas included D261 and E265 in NEK7 pairing with residues in the Helical Domain 2 (HD2) in NLRP1 and NLRP3. These interactions were limited in our simulations of NLRP3 to hydrogen bonds between D261 and K696 in the HD2 in 2 out of the 5 replicas. On the contrary, R693, R698 and S715 in NLRP1 interact with D261 and E265 in NEK7 in all the simulations run. Lastly, the interactions involving the nucleotide binding domain (NBD) of NLRP3 were not reported to have drastic effects on the overall binding to NEK7. We observed a limited number of interactions in the interface between NBD and NEK7 that included mainly the formation of salt bridges involving D290, K293 and R294 in NEK7. Apart from the specific interactions reported by Sharif and colleague (Sharif et al., 2019), we observe the formation of many other transient pairwise interactions, mainly involving charged and polar residues, in the contacting interfaces of NEK7, NLRP3 and NLRP1.

**Figure 1.**
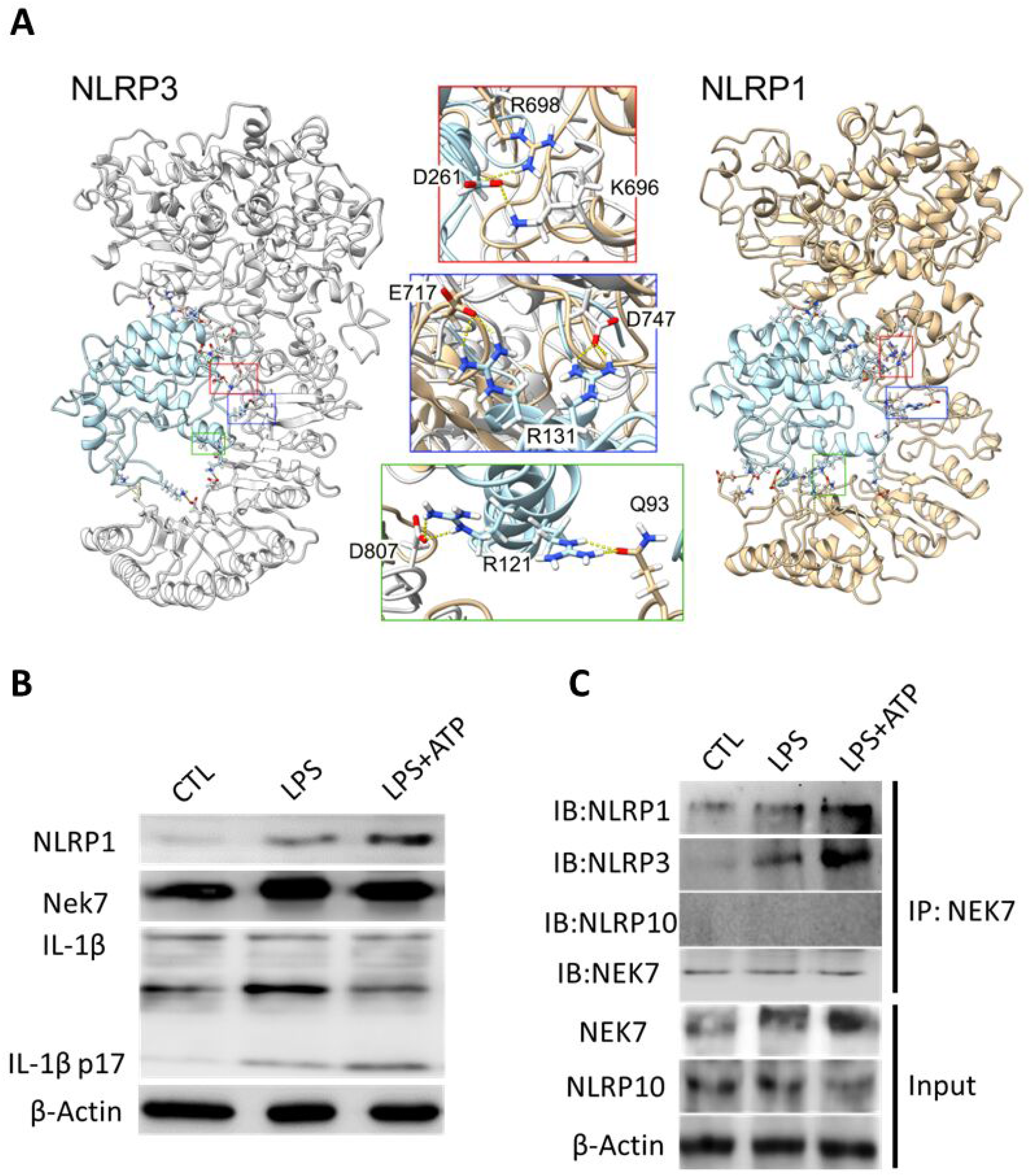
NLRP1 interacts with NEK7. (A) Overall view of the quaternary structures of the most representative cluster obtained from the MD simulations of NLRP3 (colored silver) and NLRP1 (colored tan) in complex with NEK7 (colored light blue). Three detailed views of pairwise interactions highlighted in the main text are shown with the same coloring scheme. (B) Undifferentiated THP-1 cells were stimulated with LPS (1 µg/ml, 4 h) and LPS plus ATP (5 mM, 30 min). Cell lysates were analyzed by immunoblotting with the indicated antibodies. (C) THP-1 cells were stimulated with LPS (1 µg/ml, 4 h) and LPS plus ATP (5 mM, 30 min). Cell lysates were immunoprecipitated (IP) and immunoblotted (IB) with indicated antibodies. Results are representative of three independent experiments.

To corroborate these modeling findings, we carried out biochemical studies. NLRP1 protein expression was induced in THP-1 cells by LPS and LPS+ATP and was associated with IL-1β maturation (Fig. 1B and Fig. S1). These stimulating factors did not affect NEK7 protein expression (Fig. 1B and Fig. S1). Co-immunoprecipitation studies revealed that NEK7 associated not only with NLRP3, as expected, but also with NLRP1 while failing to interact with NLRP10, which does not have LRR domain (Fig. 1C and Fig. S2). These findings suggest that the LRR domain might be important for NEK7 binding to NLRP1 or NLRP3.

LPS and ATP are well known activators of the NLRP3 inflammasome. To directly actívate the NLRP1 inflammasome, we used the DPP8/9 inhibitor, Val-boroPro (VbP), and promoted caspase-1 maturation as assessed by the generation of caspase-1 p20 fragment (Fig. 2A and Fig. S3). To determine whether NLRP1 interacts with NEK7 in response to Vbp treatment, we performed coimmunoprecipitation studies. NLRP1 associated with NEK7 in non-stimulated cells, a response that was amplified upon Vbp stimulation (Fig. 2B and Fig. S4). While ASC recruitment to NLRP1-NEK7 complex was weak at baseline, this response was increased upon Vbp stimulation, and correlated with the activation of the NLRP1 inflammasome based on caspase-1 activation (Fig. 2B and Fig. S4).

**Figure 2.**
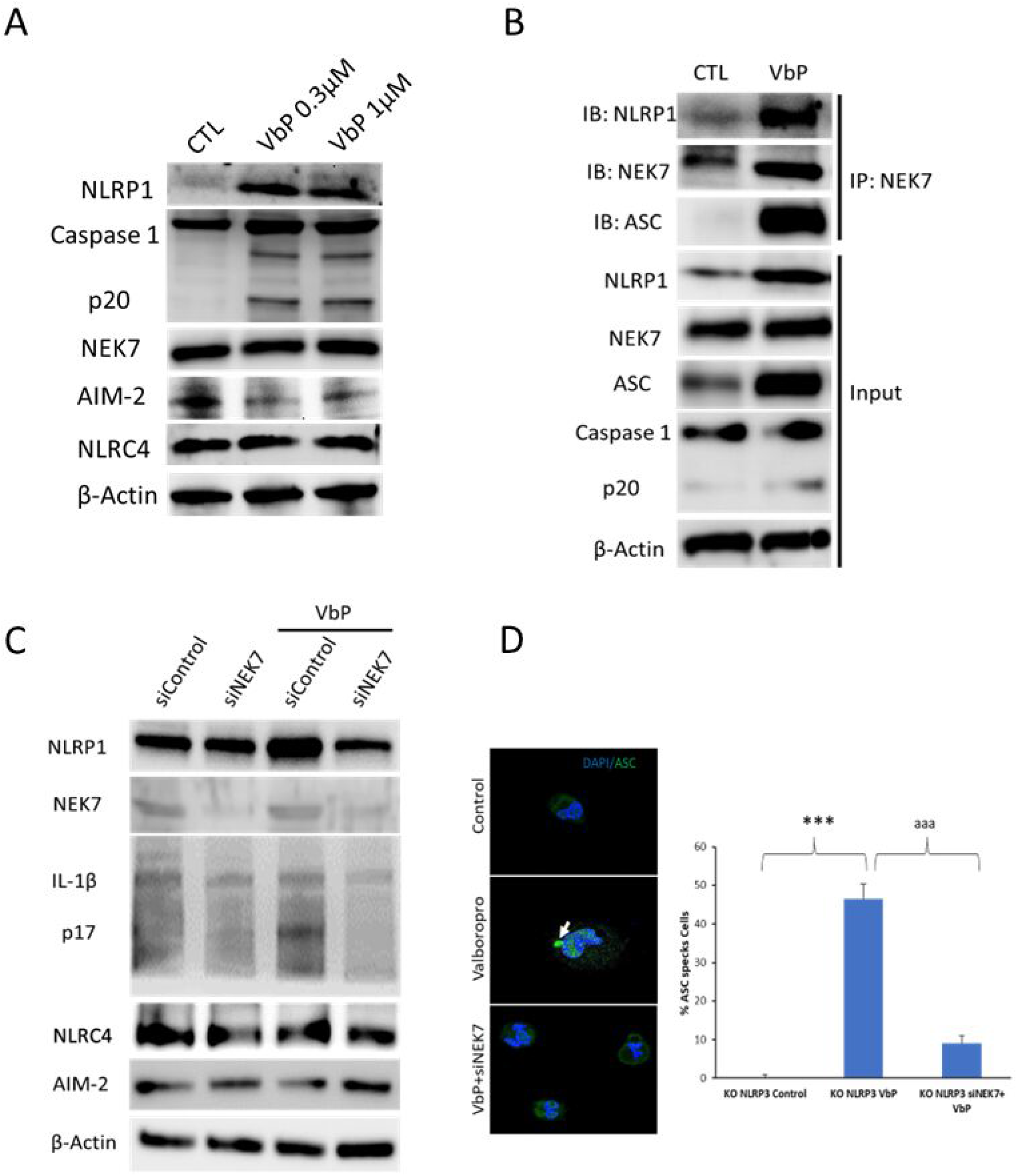
NLRP1 interaction with NEK7 is independent of NLRP3. (A) Undifferentiated NLRP3 KO THP-1 cells were stimulated with Valboropro (VbP) 0.3 and 1µM (4h). Cell lysates were analyzed by immunoblotting with the indicated antibodies. (B) Undifferentiated NLRP3 KO THP-1 cells were stimulated with Valboropro (VbP) 0.3µM (4h). Cell lysates were immunoprecipitated (IP) and immunoblotted (IB) with indicated antibodies. (C) NLRP3 KO THP-1 cells were stimulated with Valboropro (VbP) 0.3µM (4h) and was treated with control siRNA (siControl) or NEK7 siRNA (siNEK7). (D) Differentiated NLRP3 KO macrophages from THP-1 cells were stimulated with VbP 0.3µM (4h) and was treated with siNEK7. The blue signal corresponds to DAPI staining and the green signal to ASC. We show percentage of ASC specks in the cells treated. Data represent the mean ± SD of three independent experiments (***p < 0.001).

To assess the functional impact of NEK7, on NLRP1 inflammasome activation, we analyzed THP1 cells in which NEK7 was knocked down usingspecific small-interfering RNA (siRNA) (siNEK7). NEK7 levels were significantly reduced in siNEK7 compared to cells transfected with control siRNA (Fig. 2C and Fig. S5). While NLRP1 expression is not affected by siNEK7 in unstimulated cells as compared with non-targeting siRNA, it was not induced in siNEK7 cells treated with VbP (Fig. 2C and Fig. S5). Importantly, NEK7 knockdown remarkably attenuated IL-1β processing induced by VbP whereas the expression of NLRC4 and AIM-2 was unaffected (Fig. 2C and Fig. S5). Furthermore, we used confocal microscopy to monitor the consequence of NEK7 knocdown on the formation of ASC specks in response to VbP treatment. Vbp induced ASC speck formation, an outcome than significantly reduced cells treated with siNEK7 (Fig.2D). Collectively, these results suggest a role of NEK7 in the activation of NLRP3 and NLRP1 inflammasomes.

The NLRP1 and NLRP3 inflammasomes are critical components of innate immunity that promote host defense, yet their aberrant activation contributes to the pathogenesis of several inflammatory diseases (Tupik et al., 2020). While the molecular mechanism of NLRP3 inflammasome activation has been thoroughly studied those of the NLRP1 inflammasome remains poorly understood. The activation of the inflammasome is associated with decreased intracellular potassium concentrations as inhibition of K+ efflux prevents this response (Pétrilli et al., 2007). NLRP3 inflammasome activation involves the participation of many proteins, including NEK7, which is induced by various stimuli such as K+ efflux, reactive oxygen species, and NF-κB signaling (Sharif et al., 2019, Deng et al. 2019, He et al., 2016, Groß et al., 2016, Chen et al., 2019). Some of these pathways also activate the NLRP1 inflammasome (Bleda et al., 2017, Dong et al., 2020), suggesting potential interactions of NEK7 with NLRP1 (Pétrilli et al., 2007, He et al., 2016).

Our study shows evidences of NEK7 and NLRP1 interaction since inhibition of NEK7 impairs NLRP1 inflammasome activation. These findings suggest dual actions of NEK7 as a common regulator of the NLRP1 and NLRP3 inflammasome pathways. The actual model of NLRP1 inflammasome activation suggests a proteasomal degradation of the entire N-terminal NOD-LRR-ZU5 fragment to liberate the active UPA-CARD fragment, which rapidly oligomerizes to engage downstream inflammasome effectors such as ASC and pro-caspase-1 (Huang et al., 2021). Our study does not contradict this model because the LRR domain, which binds NEK7 has been proposed to have a role on the NLRP1 activation before its proteasomal degradation. For example, LRR has been shown to bind nucleic acids with high affinity, acting as a RNA sensor (Bauernfried et al., 2020). In summary, our study proposes a novel regulatory mechanism of the NLRP1 inflammasome by NEK7. Thus, NEK7 might be therapeutically targeted to prevent NLRP3 and NLRP1 inflamasomopathies.

## Material and Methods

### Reagents

Monoclonal antibodies specific for NLRP1 and NLRP3 were purchased from Novus Biologicals (Colorado, USA). Recombinant anti-NEK7 monoclonal antibody was acquired from Abcam (Cambridge, UK). Similarly, anti-AIM-2, Caspase 1 and IL-1β were obtained from Cell Signaling Technology (Beverly, MA, USA). Finally, NLRC4 was obtained from Merck (Darmstadt, Germany). Goat Anti-Rabbit IgG H&L (HRP), goat Anti-Mouse IgG, H&L Chain Specific Peroxidase Conjugate, lipopolysaccharides from Escherichia coli (LPS), ATP and Poly(deoxyadenylic-thymidylic) acid sodium salt (Poly dA-dT) were obtained from Merck (Darmstadt, Germany). Val-boroPro - Calbiochem 5314650001 was obtained from Merck (Darmstadt, Germany). A cocktail of protease inhibitors (Complete™ Protease Inhibitor Cocktail) was purchased from Boehringer Mannheim (Indianapolis, IN). The Immun Star HRP substrate kit was obtained from Bio-Rad Laboratories Inc. (Hercules, CA). Finally, siRNAS of control and NEK7 were obtained from ThermoFisher, Waltham, MA, USA.

### Cell culture and treatments

THP-1 cells were cultured in RPMI-1640 supplemented with 2 mM l-glutamine, 10% FBS, and Pen/Strep (100 U/mL) and incubated at 37 °C, 5% CO_2_. Cells were primed with 200 ng mL-1 ultrapure LPS for 4 h, followed by stimulation with 5 mM ATP (30 min). Valboropro were used 0.3 and 1µM and Poly(dA-dT) DNA was transfected using Lipofectamine 2000 at a concentration of 1 µg/ml.

### All-atom MD simulations of NLRP3-NEK7 and NLRP1-NEK7 complexes

The Cryo-EM structure of the NATCH and LRR domains of NLRP3 in complex with the C-terminal lobe of NEK7 (PDB 6NPY) (3) was used as a template for generating the model of the equivalent structural domains of NLRP1 (residues 227-990) with Modeller v10.1 (ref). We also used Modeller to fill in the short fragments of NLRP3 and NEK7 not resolved in the Cryo-EM structure. The structures of the NLRP1 and NLRP3 complexes were solvated in a 1.0 nm edge truncated octahedron of TIP3P waters, and counterions were added until neutralization.

All the MD simulations have been run locally with Gromacs v2020.5 (reference) using amber ff03 (reference). The initial energy minimization protocol comprised 50000 steps of steepest descent, followed by a 250 ps heating process (time step 1.0 fs) up to 303 K. A very short equilibration process with a time step of 1.0 fs and 303 K temperature was run for 100 ps, followed by a 100 ns equilibration where the time step was increased to 2.0 fs. A production simulation and 5 replicate trajectories were run for 100 ns each. Long range electrostatic calculations were carried out with the Particle Mesh Ewald method and a cutoff value of 1.2 nm. Temperature and pressure coupling were performed using the Nose-Hoover and Parrinello-Rahman ensembles, respectively. A total of 1000 frames were saved in each simulation run. The calculation of RMSD and number of hydrogen bonds and clustering analysis in the MD trajectories were also carried out with Gromacs functions rmsd, hbond and cluster, respectively.

### Immunoblotting

Western blotting was performed using standard methods. After protein transfer, the membrane was incubated with various primary antibodies diluted 1:1000, and then with the corresponding secondary antibodies coupled to horseradish peroxidase at a 1:10000 dilution. Specific protein complexes were identified using the Immun Star HRP substrate kit (Biorad Laboratories Inc., Hercules, CA, USA).

### Coimmunoprecipitation

Cells were cultured on T75 flasks until 90% confluence, then scrapped off with ice cold PBS 1x and centrifuged at 1.000g for 5’. Cell pellets were lysed using native lysis buffer (ab156035, Abcam, Cambridge, UK) with 1% PMSF and briefly sonicated for homogenization. Cellular solution was centrifuged at 16.000g for 2’ to collect proteins from the supernatant. Protein concentration was quantified by BCA Pierce Assay (23225, ThermoFisher, Waltham, MA, USA). For crosslinking preparation, samples were diluted to 1μg/μl protein using the lysis buffer and then the antibody of interest was added at a rate of 1:50. Samples were incubated overnight at 4ºC in agitation.

For protein isolation, Pierce protein A magnetic beads were used (8846, ThermoFisher, Waltham, MA, USA). Magnetic beads were vortexed and wash 3 times with PBS Tween 0.1%, using the magnetic rack (6-Tube SureBeads™ Magnetic Rack #1614916 BIORAD Hercules, CA, USA) to magnetized them. Then previous samples were mixed with the beads and incubated 1 hour in agitation at 4ºC. After the incubation, samples were placed on the magnetic rack on ice. Supernatant was kept as a control of the immunoprecipitation. Beads were wahsed 3 times with cold PBS-T using the magnetic rack. Samples were eluted using Laemmli buffer and heated for 10’ at 70ºC. Finally, released beads were retrieved using the magnetic rack. Isolated proteins were used for standard Western blot assays.

#### NEK7 knockdown in THP-1 cells and derived macrophages

Cells were seeded on 6-wells plates until 75% confluence in 2ml DMEM high glucose medium (Cat. 10566016) supplemented with 10% FBS and 1% antibiotics. Transfection was performed according the lipofectamine RNAiMAX reagent (Cat. 13778-075) protocol. Briefly, the siRNA-lipid complex was prepared in DMEM medium with 3% lipofectamine and 30 pmol NEK7 siRNA (Cat. AM51331), and incubated for 5 minutes at RT to form the silencing complex. Then, 250 µl of the siRNA-lipid complex were added in each well. After 72h, cells were treated and analysed for the different conditions. Every reagent including DMEM medium was purchased from ThermoFisher (Waltham, MA, USA).

#### Statistical análisis

Data are expressed as mean ± SD. Statistical analysis was performed using unpaired two-tailed Student’s t test or Mann-Whitney test by GraphPad Prism. A p value less than 0.05 was considered statistically significant

## Acknowledgments

We thank Dr. Veit Hornung (Ludwig Maximilian University, Munich) and Pablo Pelegrin (Biomedical Research Institute of Murcia) for providing the NLRP3 KO THP-1 cells. This study was supported by PI21/01656 grant, Instituto de Salud Carlos III, Spain.

## Conflict of interests

The authors declare that the research was conducted in the absence of any commercial or financial relationships that could be construed as a potential competing interest.

